# T505A variant of ETV5 promotes proliferation of precursor B cells in a mouse model of acute lymphoblastic leukemia

**DOI:** 10.1101/2024.12.19.629436

**Authors:** Joshua Yi, Michael Wu, Michaela L. Dowling, Allanna C. E. MacKenzie, James Iansavitchous, Rodney P. DeKoter

## Abstract

Precursor B cell acute lymphoblastic leukemia (pre-B-ALL) arises as a result of precursor B cells acquiring driver mutations that lead to arrested differentiation and increased proliferation. Identification of driver mutations and understanding their biological function is critical to understanding pre-B-ALL development and advancing disease treatment. Using a mouse model of pre-B-ALL driven by deletion of genes encoding the related E26-transformation-specific (ETS) transcription factors PU.1 and Spi-B, we performed whole exome sequencing to identify secondary driver mutations. We identified recurrent variants in E26 transformation-specific transcription variant 5 (ETV5) resulting in R392P, V444I, and T505A amino acid changes. We found that the R392P and V444I variants altered the ability of ETV5 to bind to DNA using electrophoretic mobility shift assay. R392P and V444I variants did not activate a Dual-Specificity-Phosphatase 6 (DUSP6) reporter. In contrast, T505A ETV5 could interact with DNA and activate the DUSP6 promoter. To determine biological function, we forced expression of wild type, R392P, V444I, or T505A ETV5 in an interleukin-7-dependent pre-B cell line. Proliferation and apoptosis assays showed that T505A ETV5 conferred a proliferative advantage to pre-B cells. RNA sequencing showed that expression of ETV5 variants significantly altered gene expression in cultured cells. Through gene set enrichment analysis, T505A was suggested to downregulate the p53 pathway and the anti-proliferative protein, B cell translocation gene 2 (encoded by *Btg2*). In summary, these data suggest that ETV5 mutations play a role in pre-B-ALL by affecting proliferation and cell survival.

## Introduction

Precursor B cell acute lymphoblastic leukemia (pre-B-ALL) is the most common cancer in young children (Hunger & Mullighan, 2015). Remarkably, the cure rate for pediatric pre-B-ALL has risen to >90%; yet, despite promising prognoses, B-ALL remains a leading cause of pediatric cancer-related deaths (Iacobucci & Mullighan, 2017; Mullighan, 2019). Pre-B-ALL arises as a result of precursor B cells acquiring driver mutations that lead to differentiation arrest and subsequent proliferation of leukemic blasts in the bone marrow (Mullighan, 2012). Identifying these driver mutations is critical in understanding B-ALL development and advancing leukemia treatment.

We previously identified recurrent driver mutations in *Jak1* and *Jak3* in the Mb1-Cre*Δ*PB mouse model of pre-B-ALL (Lim *et al*, 2020; Batista *et al*, 2018). The Mb1-Cre*Δ*PB mouse is homozygous null for the E26-transformation-specific (ETS) transcription factor Spi-B (encoded by *Spib*) and deletes the related ETS transcription factor PU.1 (encoded by *Spi1*) during B cell development under the control of Mb1-(encoded by *Cd79a*) Cre (Batista *et al*, 2017). Leukemias arise in 100% of Mb1-Cre*Δ*PB mice with a median time to euthanasia of 18 weeks (Batista *et al*, 2018). Treatment of Mb1-Cre*Δ*PB mice with the Janus Kinase inhibitor ruxolitinib significantly delayed leukemia progression (Lim *et al*, 2020). Whole-exome sequencing showed that leukemias that progressed in ruxolitinib-treated mice had decreased *Jak1* and *Jak3* mutation frequencies and an increased diversity of mutational signatures (Lim *et al*, 2020). However, it was not clear what mutations promoted leukemia progression in ruxolitinib-treated mice.

E26 transformation-specific transcription variant 5 (ETV5), also known as ETS-related molecule (ERM), belongs to the polyomavirus enhancer activator 3 (PEA3) subfamily of ETS transcription factors, which consists of ETV1, ETV4, and ETV5 (Monté *et al*, 1994). PEA3 transcription factors are characterized by an N- and C-terminal acid transactivation domain (TAD), a central negative regulatory domain (NRD), and an ETS DNA binding domain that binds to the purine-rich GGAA DNA sequence (Currie *et al*, 2017; Laget *et al*, 1996). PEA3 subfamily members are well-established oncogenic factors with a wide range of cancer-promoting functions (Currie *et al*, 2017). Specifically, ETV5 regulates multiple biological processes in cancer, such as the cell cycle, apoptosis, angiogenesis, epithelial-mesenchymal transition, and drug resistance (Wei *et al*, 2023). ETV5 has been implicated in multiple solid tumour types including but not limited to bladder, breast, brain, colorectal, gastric, lung, ovarian, thyroid, and prostate cancer (Wei *et al*, 2023).

Several pieces of emerging evidence point to ETV5 as an important player in the development of pre-B-ALL. Firstly, *Etv5*, alongside other genes, such as *Dusp6* and *Spry2*, was shown to be critical for progression of pre-B-ALL in mice (Shojaee *et al*, 2015). ETV5 was found to induce oncogenic transformation through negative regulation of Erk signaling (Shojaee *et al*, 2015). Knockout of *Etv5* in BCR-ABL-transgenic mice was shown to prevent negative regulation of Erk, resulting in the promotion of an anti-pre-B-ALL profile including increased expression of p53 tumor suppressors and decreased expression of pre-B cell genes. Moreover, *Etv5*^-/-^ tumors were found to be negatively selected from the pre-leukemic repertoire, indicating that ETV5 and Erk negative regulation is critical for the development of pre-B-ALL (Shojaee *et al*, 2015). ETV5 was also found to upregulate the aPKCλ/ι-SATB2 (atypical protein kinase λ/ι-AT-rich sequence-binding protein 2) signaling cascade, which is required for oncogenic transformation in BCR-ABL+ leukemias (Nayak *et al*, 2019). Using BCR-ABL transgenic mice, ectopic expression of ETV5 was shown to upregulate expression of the transcription factor SATB2, leading to increased proliferation and malignant transformation of progenitor B cells (Nayak *et al*, 2019). Taken together, this suggests a potential role of ETV5 in pre-B-ALL; however, the underlying mutations contributing to ETV5-mediated leukemogenesis remain relatively unexplored.

In this study, using whole exome sequencing data we discovered that *Etv5* variants were frequent in leukemias from ruxolitinib-treated Mb1-Cre*Δ*PB mice. We investigated the function of R392P, V444I, and T505A ETV5 variants since mutations at these positions have also been observed in human cancers. We found that the R392P and V444I variants altered the ability of ETV5 to bind to DNA and abrogated activation of the Dual-Specificity-Phosphatase 6 (DUSP6) promoter. In contrast, T505A ETV5 could bind DNA and activate the DUSP6 promoter. To determine biological function, we forced expression of wild type, R392P, V444I, or T505A ETV5 in the SeptMBr interleukin-7-dependent pre-B cell line. Cell counting and proliferation assays demonstrated that T505A ETV5 conferred a proliferative advantage to pre-B cells. RNA sequencing showed that expression of ETV5 variants significantly altered gene expression in cultured cells. Through gene set enrichment analysis, T505A was suggested to downregulate the p53 pathway and the anti-proliferative protein, B cell translocation gene 2 (encoded by *Btg2*). In summary, these data suggest that ETV5 mutations play a role in pre-B-ALL by affecting proliferation and cell survival.

## Results

### Identification of Etv5 genetic variants in Mb1-Cre⊗PB leukemias

The Janus Kinase inhibitor ruxolitinib delays leukemia progression in the Mb1-Cre**⊗**PB mouse model and reduces *Jak1/Jak3* mutation frequencies (Lim *et al*, 2020). To determine if ruxolitinib treatment affected other driver mutation frequencies, whole-exome sequencing (WES) data from 11 leukemias and matched control tail DNA samples (5 from control mice, 7 from Ruxolitinib-treated mice) was analyzed using Strelka and Varscan variant callers (**Table 1, Fig S1A, Table S2**). Varscan called genetic variants from WES data with relaxed specificity compared to Strelka, calling an average of 1061 variants compared to Strelka calling an average of 130 variants at 10% or greater variant allele frequency (VAF) (**Fig S1B, Table S2, variant lists in supplemental data**). Since Strelka’s high stringency limited the detection of variants, Varscan results were used for further analysis. Mutational signature analysis performed using DeconstructSigs (Rosenthal *et al*, 2016) showed that C->A and C->T variants were frequent in Mb1-Cre**⊗**PB leukemias (**Fig S1C**). To determine if C->A transversions or C->T transitions occurred at distinct variant allele frequencies, we placed frequencies into bins. We found that in low VAF bins (0.15-0.4) the ratio of C->A/C->T mutations was significantly higher than the ratio in high VAF bins (0.4-0.65) (**Fig S1D**). This result suggests distinct mutational processes occurring early in clonal evolution (high VAF) compared to late in clonal evolution (low VAF).

**Table 1.** Characterization of Etv5 variants identified in a mouse model of precursor B Cell Acute Lymphoblastic Leukemia. Whole-exome sequencing was performed on Mb1-CreDPB mouse leukemias (indicated by number) with or without ruxolitinib treatment. ETV5 variants were characterized by mutation type, variant allele frequency, nucleotide change, and whether a human variant at the same position was identified in the COSMIC catalog of somatic mutations.

| **Mouse** | **Ruxolitinib** | **Variant** | **Mutation type** | **Variant Allele Frequency** | **Nucleotide Change** | **Human Equivalent** |
| --- | --- | --- | --- | --- | --- | --- |
| 224 | Yes | T505A | Missense | .273 | T->C | A505V |
| 224 | Yes | M496K | Missense | .253 | A->T | none |
| 226 | Yes | T505A | Missense | .250 | T->C | A505V |
| 226 | Yes | M496K | Missense | .118 | A->T | None |
| 261 | Yes | M496K | Missense | .182 | A->T | None |
| 307 | Yes | V444I | Missense | .103 | C->T | V444I |
| 310 | Yes | M496K | Missense | .375 | A->T | none |
| 310 | Yes | T505A | Missense | .300 | A->T | A505V |
| 486 | Yes | T505A | Missense | .364 | T->C | A505V |
| 486 | Yes | M496K | Missense | .307 | A->T | none |
| 486 | Yes | D6Y | Missense | .125 | C->A | D6H |
| 262 | No | T505A | Missense | .222 | T->C | A505V |
| 484 | No | R392P | Missense | .100 | C->G | R392Q |

Next, we identified genes in which variants with VAF > 10% occurred one or more times in the 11 leukemias tested. First, intramouse duplicates were removed. Second, genes were prioritized that had 7 or more variants (**Table S3**). Third, genes were prioritized that appear in the COSMIC Cancer Gene Census (Tate *et al*, 2019). *Etv5* was of interest since it encodes a member of the E26-transformation-specific (ETS) transcription factor family that is highly related to PU.1 and Spi-B (Sharrocks, 2001). In total, 8/11 leukemias had at least one *Etv5* variant, and 4/11 leukemias had more than one *Etv5* variant (**Table 1**). There were five different single-nucleotide missense variants identified in *Etv5*, D6Y (1/11 leukemias), T505A (5/11 leukemias), M496K (5/11 leukemias), V444I (1/11 leukemias) and R392P (1/11 leukemias). Interestingly, 6/7 ruxolitinib-treated mice had *Etv5* variants, while 2/4 control mice had an *Etv5* variant (**Table 1, Table S2**), suggesting that *Etv5* mutation frequencies may have been affected by ruxolitinib treatment.

Murine and human ETV5 proteins have 96% sequence identity and contain 510 amino acids (Monté *et al*, 1994). D6 is located at the N-terminal region of the protein. R392 and V444 are located within the ETS DNA-binding domain, while M496 and T505 are located in the C-terminal region of the protein (**Fig 1A, B**). The ETS domain of ETV5 is evolutionarily conserved, and R392 and V444 are identical between multiple species. Interestingly, T505 occurs as an A in most species, with three strains of mice including C57Bl/6 as outliers (**Fig 1C**). Therefore, the T505A variant changes this amino acid into the alanine that occurs in most species including human (**Fig 1C**). The COSMIC cancer gene census records six human D6H mutations, two R392Q mutations, one V444I mutation, and four A505V mutations, but no M496 mutations (Tate *et al*, 2019). Based on these findings, R392P, V444I, and T505A ETV5 variants were prioritized for further study.

**Figure 1.**
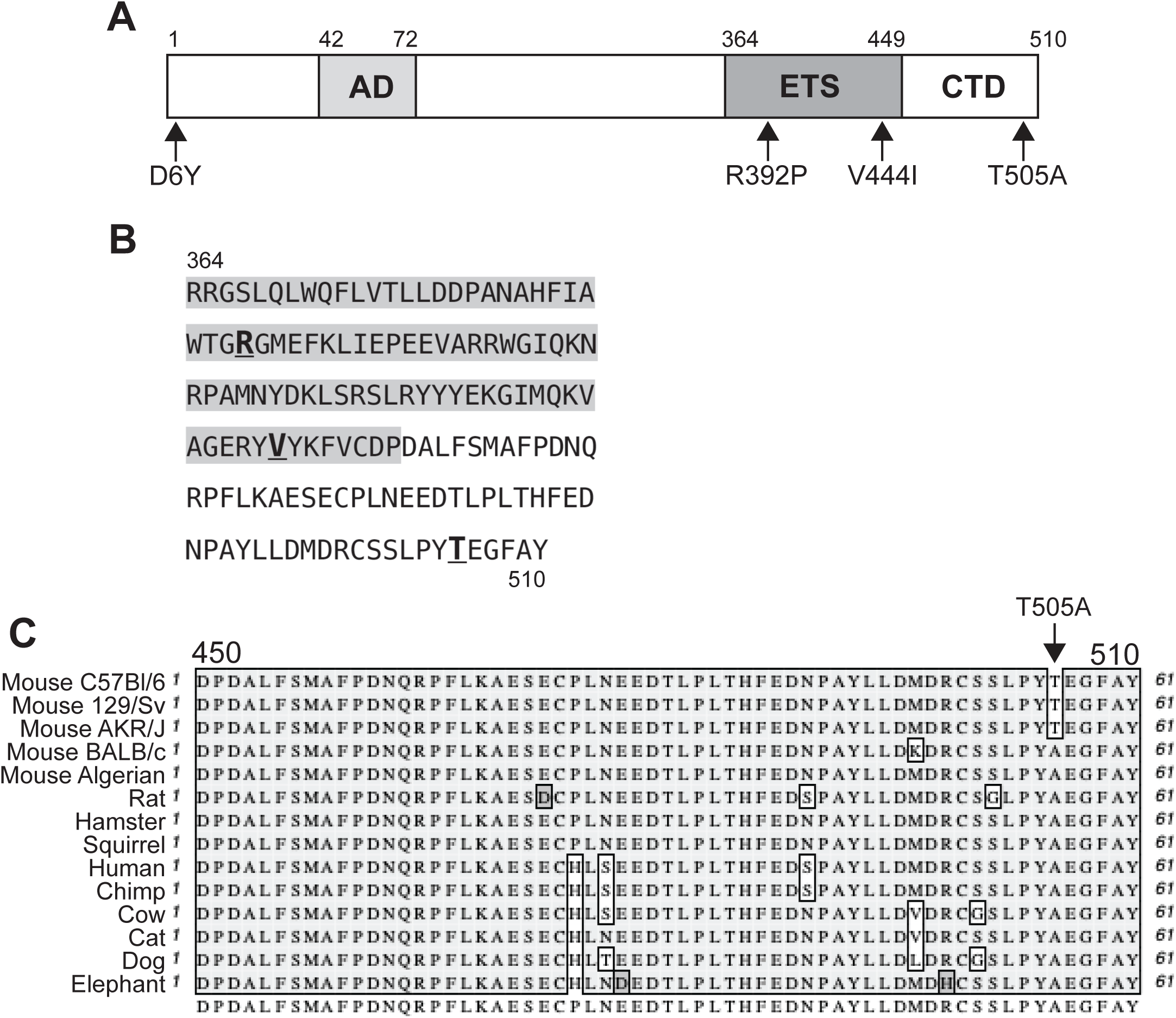
Location of ETV5 amino acid variants. **A)** Schematic of ETV5 Protein Structure. ETV5 protein (510 amino acids) contains an N-terminal activation domain (AD), a DNA binding domain (ETS), and a C-terminal domain (CTD) at the indicated positions. **B)** Amino acid sequence of ETV5 from positions 364-510. Locations of R392P, V444I, and T505A variants are bolded and underlined. **C)** Multispecies amino acid sequence alignment of ETV5 positions 450-510. Species are indicated on the left. Position of T505 is indicated with an arrowhead.

### Electrophoretic mobility shift assay and Luciferase assay of ETV5 proteins

To assess the effects of the R392P, V444I, and T505A ETV5 variants on DNA binding ability, mutants were generated in mouse *Etv5* cDNA ligated into the pBlueScript KS+ vector. *In vitro* translated (IVT) proteins were detectable by immunoblot using polyclonal anti-ETV5 Ab (**Fig 2A**). IVT products containing wild type, R392P, V444I, and T505A ETV5 exhibited two distinct bands at ∼45 and ∼70 kDa, which were absent in the empty pBlueScript KS+ lane (**Fig 2A**). Bands at ∼70 kDa corresponded to full-length ETV5, while bands at ∼45 kDa likely represented truncated ETV5 initiating at nucleotide position 754 (**Table S4**).

**Figure 2.**
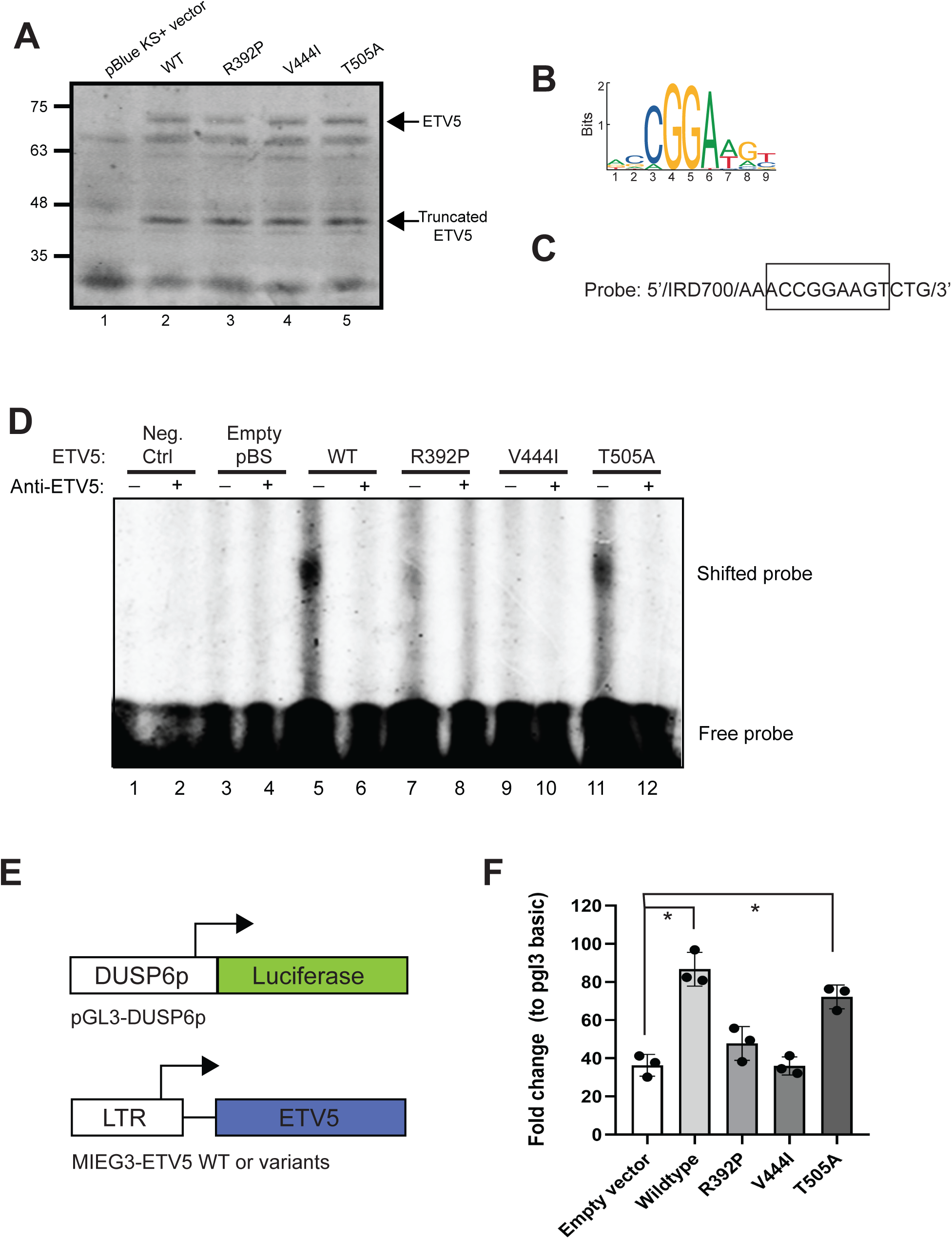
R392P and V444I ETV5 variants have altered DNA binding activity. **A)** Immunoblot of *in vitro* translated wild type (WT), R392P, V444I, and T505A ETV5 proteins. Molecular weight (kDa) is indicated on the left. Negative control is pBlue KS+ vector. Full-length and truncated ETV5 are indicated with arrowheads. **B)** Consensus ETV5 DNA binding logo derived from ChIP-seq data. **C)** DNA sequence of oligonucleotide probe for ETV5 based on consensus DNA binding logo. **D)** Electrophoretic mobility shift analysis (EMSA) using *in vitro* translated ETV5 proteins from (A) and oligonucleotide probes from (B). Anti-ETV5 indicates lanes in which binding reactions were incubated with anti-ETV5 antibody. Location of shifted and free probe is indicated on the right. **E)** Schematic of DUSP6 promoter luciferase reporter vector (top) and MIEG-ETV5 expression vector (bottom). **F)** Activation of DUSP6 promoter by ETV5. DUSP6 promoter luciferase reporter was activated in HeLa cells by wild type (wildtype) and T505A ETV5, but not by R392P or V444I ETV5. *p<0.05 by one-way ANOVA and Tukey’s test.

ETV5 was recently reported to preferentially interact with the core sequence CCGGAA (Zhang *et al*, 2017) (**Fig 2B**). Therefore, we performed electrophoretic mobility shift assay (EMSA) using IVT proteins and an oligonucleotide probe containing the ETV5 consensus sequence shown in **Fig 2C**. No shifted bands were observed in negative controls (probe alone and empty pBlueScript KS+, **Fig 2C lanes 1 and 2**). A clearly shifted band was observed for wild type ETV5 (**Fig 2D, lane 5**). The shifted band was not visible upon addition of anti-ETV5 polyclonal Abs (**Fig 2D, lane 6**). Wild type ETV5 protein did not interact with mutated probe (**Fig S2**). R392P ETV5 exhibited a less intense shift that was eliminated with the addition of antibody (**Fig 2D, lanes 7 and 8**). V444I ETV5 exhibited no detectable binding with or without anti-ETV5 polyclonal Abs (**Fig 2D, lanes 9 and 10**). Finally, T505A ETV5 exhibited a shift that was similar to wild type protein (**Fig 2D, lanes 11 and 12**). R392P, V444I, and T505A ETV5 proteins did not interact with the mutated probe (**Fig S2**). These results suggested that the R392P and V444I mutations altered DNA-binding ability, while the T505A mutation had no detectable effect on DNA-binding ability. Next, the effect of R392P, V444I, and T505A ETV5 variants were tested on target gene activation. The Dual-Specificity Phosphatase-6 (DUSP6) promoter has been reported to be a direct target of activation by ETV5 (Shojaee *et al*, 2015; Znosko *et al*, 2010). We cloned a 782 bp fragment of the mouse *Etv5* promoter into the pGL3-basic luciferase reporter and determined if wild type or variant ETV5 could activate this promoter in a transient transfection experiment (**Fig 2E**). The DUSP6 promoter showed activity above background upon transfection into HeLa cells or 38B9 pre-B cells (**Fig S3A, B**). Wild type and T505A, but not R392P or V444I ETV5 trans-activated the DUSP6 promoter in HeLa cells (**Fig 2F**). Since the T505A variant occurs near a C-terminal trans-activation domain of ETV5 (Laget *et al*, 1996) we also tested the ability of a wild type and T505A C-terminal fragment of ETV5 to trans-activate the DUSP6 promoter. Both wild type and T505A C-terminal ETV5 trans-activated the DUSP6 promoter (**Fig S3C, D**). Overall, these results indicated that wild type and T505A ETV5 trans-activated the *Dusp6* promoter, while R392P and V444I ETV5 failed to activate the *Dusp6* promoter.

### T505A ETV5 Promotes Proliferation of Mouse Pre-B cells

To study the effect of ETV5 mutations on proliferation and survival, the pre-B cell line SeptMBr was transduced with MIEG3 retroviral vectors encoding wild type, R392P, V444I, or T505A ETV5. SeptMBr cells are interleukin-7-dependent and are genetically deleted for genes encoding PU.1 and Spi-B, as were the leukemias in which *Etv5* mutations were discovered, providing an appropriate context to test ETV5 variant function (Sams *et al*, 2024). Transduced cells were enriched using flow cytometric cell sorting (**Fig S4A**). RT-qPCR demonstrated that all ETV5-infected cell lines expressed higher levels of *Etv5* mRNA transcripts compared to the MIEG3 control (**Fig S4B**). Overexpression of ETV5 could not be reproducibly demonstrated using immunoblot because SeptMBr cells expressed endogenous ETV5 (**Fig S4C**).

To assess whether forced expression of ETV5 variants affects cell proliferation, transduced SeptMBr cells were cultured for 48 hr and cell counts were obtained using flow cytometry. R392P-transduced cells showed an increase in cell counts compared to MIEG3 (1.35-fold), wild type (1.35-fold), and V444I (1.25-fold); however, these differences were not statistically significant (**Fig 3A**). V444I-transduced cells displayed similar cell count numbers compared to MIEG3 and wild type controls (**Fig 3A**). Interestingly, T505A ETV5-transduced cells exhibited a significant increase in cell counts compared to MIEG3 (1.85-fold), wild type (1.85-fold), R392P (1.37-fold), and V444I (1.71-fold) (**Fig 3A**). Overall, these results suggested that forced expression of T505A ETV5 increased cell proliferation.

**Figure 3.**
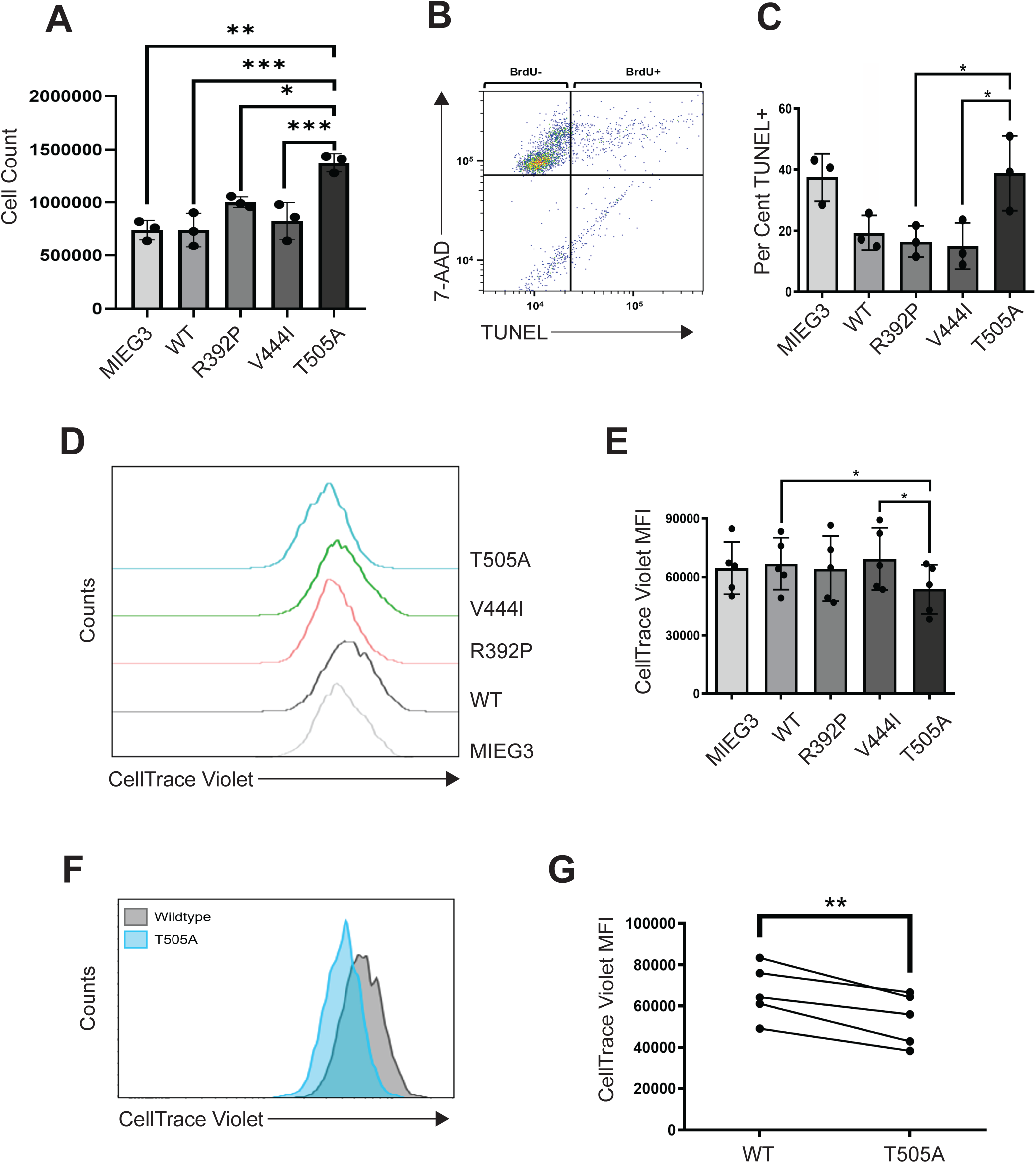
T505A ETV5 induces proliferation in SeptMBr pre-B cells. **A)** Interleukin-7-dependent SeptMBr pre-B cells were transduced with the MIEG-ETV5 retroviral vectors indicated on the x-axis and enriched by cell sorting. Y-axis indicates cell counts 48 hr after 100,000 cells were placed in culture. Each dot represents one independent biological replicate experiment. **B)** Representative flow cytometry plot for analysis of apoptosis. Y-axis indicates 7-AAD staining, x-axis indicates BrdU staining (TUNEL assay). **C)** Quantification of 7AAD+ TUNEL+ cells (upper right quadrant of panel B) from 3 biological replicate experiments. **D)** Representative histogram of flow cytometric analysis of CellTrace Violet dilution. **E)** Quantification of CellTrace Violet mean fluorescence intensity (MFI) for five independent biological replicate experiments. **F)** Representative histogram of flow cytometric analysis of SeptMBr cells transduced with wild type (wildtype) and T505A ETV5. **G)** Quantification of CellTrace Violet mean fluorescence intensity (MFI) for five independent biological replicate experiments. *p<0.05, **p<0.01, ***p<0.001 by one-way ANOVA and Tukey’s test (panels A, C, E) and by paired t-test (panel G).

To determine if increased cell numbers were due to increased survival, a deoxynucleotidyl transferase biotin-dUTP nick end labeling (TUNEL) apoptosis assay was performed. SeptMBr cells were cultured for 48 h and apoptosis was measured as a proportion of TUNEL+ cells (**Fig 3B**). The wild type, R392P, and V444I groups showed a 2-fold decrease in TUNEL+ cells compared to the MIEG3 control, suggesting enhanced survival in SeptMBr cells (**Fig 3C**). In contrast, no differences in TUNEL+ cells were observed between T505A and the MIEG3 control (**Fig 3C**). Furthermore, when compared to the wild type control, the T505A group exhibited a 2-fold increase in apoptotic cells (**Fig 3C**). In summary, these findings indicate that cells transduced with T505A ETV5 had reduced survival.

To assess whether the increased cell numbers were a consequence of increased proliferation, transduced SeptMBr cells were cultured for 48 h and proliferation was measured by dilution of CellTrace Violet dye. T505A ETV5-transduced cells exhibited a decrease in mean fluorescence intensity compared to all other groups: empty MIEG3 (−1.2-fold), wildtype (−1.2-fold), R392P (−1.2-fold), and V444I (−1.3-fold) (**Fig 3D, 3E**). Furthermore, a direct comparison between T505A and wild type ETV5-transduced cells demonstrated that the T505A group had significantly higher levels of proliferation (**Fig 3F, 3G**). Taken together, these findings suggest that the increase in T505A ETV5-transduced cell numbers may be attributed to a proliferative advantage that outpaces the rate of apoptosis.

### R392P, V444I, and T505A ETV5 induce changes in gene expression relative to wild type ETV5

To determine differences in gene expression, RNA was prepared from the MIEG3 control, wild type, R392P, V444I, and T505A ETV5-transduced SeptMBr cells, and RNA-sequencing was performed. Count tables for annotated mRNA transcripts were determined using Salmon (Patro *et al*, 2017). Principal component analysis on Log_2_-transformed count data showed that each group clustered into a distinct group (**Fig 4A**). DEseq2 (Love *et al*, 2014) was used to identify differentially expressed genes (log2FoldChange > 0.5 and padj < 0.05) between ETV5-transduced groups relative to the MIEG3 control. Surprisingly, all ETV5 mutant groups had large numbers of differentially expressed genes compared to the wild type control (**Fig 4B**). T505A ETV5-transduced cells exhibited the largest number of differentially expressed genes (753 genes), followed by R392P (718 genes), and V444I (711 genes) (**Fig 4B, gene lists in supplemental data**). Hierarchical clustering analysis showed that the identities of differentially expressed genes were quite distinct between R392P, V444I, and T505A groups relative to either MIEG3 control or wild type ETV5-transduced cells (**Fig 4C**). Genes that were up- or down-regulated in SeptMBr cells by wild type ETV5 showed limited overlap with genes affected by R392P, V444I, or T505A variants of ETV5. In other words, the genes regulated by the ETV5 variants were mostly distinct from those regulated by wild type ETV5 (**Fig. 4D-F**). Only 42 up- or down-regulated genes were in common between all groups (**Supplemental Fig 5**). In summary, forced expression of R392P, V444I, and T505A ETV5 in SeptMBr cells induced unique patterns of changes in gene expression, despite the observation that R392P and V444I ETV5 had altered DNA-binding ability.

**Figure 4.**
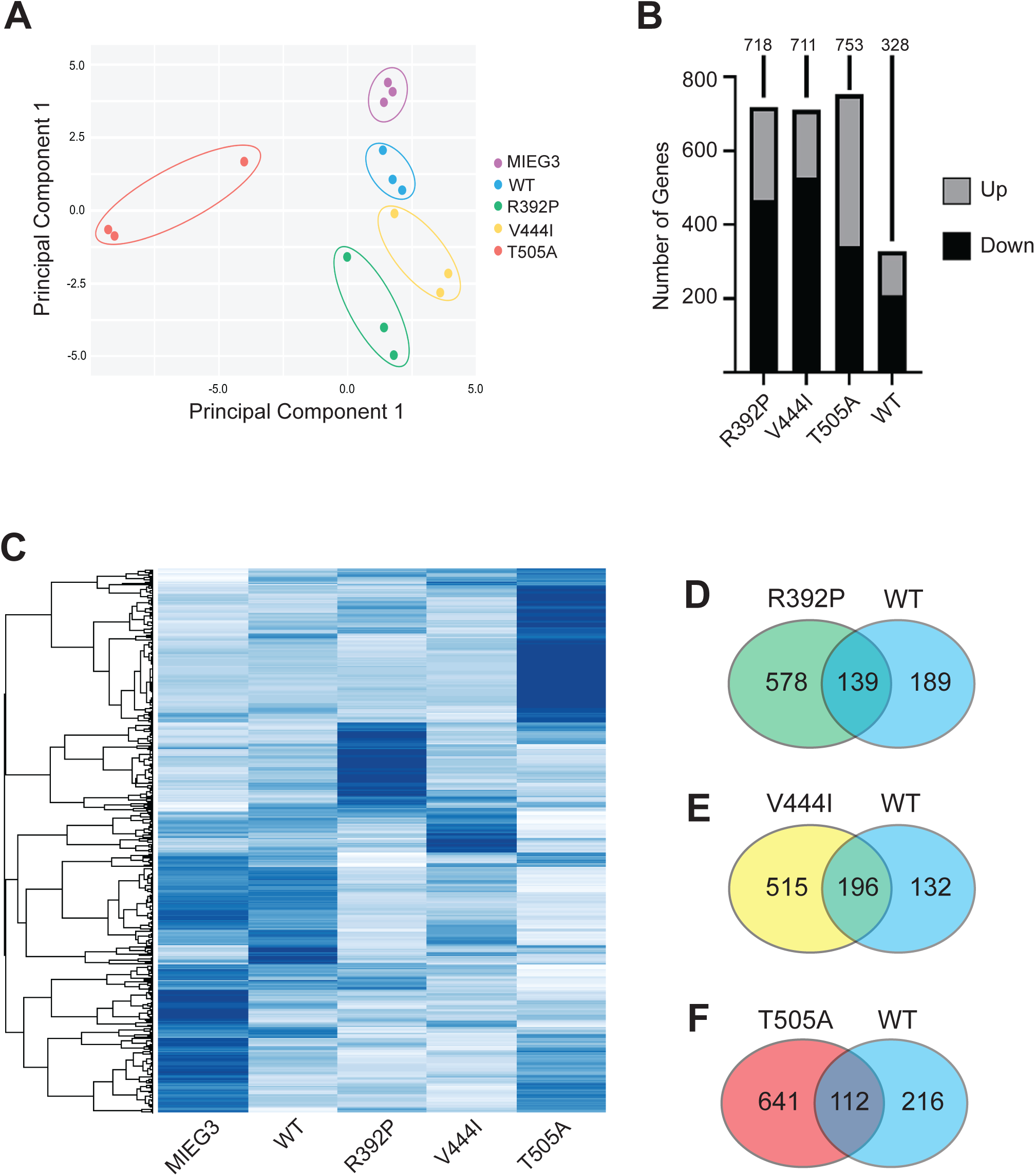
ETV5 variants induces changes in gene expression in SeptMBr pre-B cells. **A)** Principal Component Analysis (PCA) of RNA-sequencing and gene expression analysis performed on SeptMBr pre-B cells transduced with the MIEG3-ETV5 vectors indicated on the right. Analysis was performed using RNA prepared from three independent biological replicate experiments. **B)** Changes in gene expression relative to MIEG3-transduced control SeptMBr pre-B cells. The number of up-(gray) or down-(black) regulated genes is indicated on the y-axis. The total number of up- and down-regulated genes is indicated over each bar. **C)** Unsupervised clustering analysis of gene expression in SeptMBr pre-B cells transduced with retroviral vectors indicated on the x-axis. Clustering is indicated on the y-axis. **D-F)** Venn diagrams indicating the number of up- and down-regulated genes in common between SeptMBr cells retrovirally transduced with the indicated ETV5 variants.

### T505A ETV5 alters gene expression affecting survival and proliferation

In order to identify biological pathways affected by expression of ETV5 variants, gene set enrichment analysis (Subramanian *et al*, 2005) was performed with the MIEG3 control used as the reference. All variant groups affected more pathways than the wild type control (**Fig 5A**). Notably, T505A ETV5 was observed to have a significant negative enrichment in the p53 pathway (FDR = 0.001), which functions to regulate the cell cycle and maintain genome stability (Wang *et al*, 2023) (**Fig 5B**). Heatmap analysis of the top ranked genes, demonstrated that *Btg2* was the top ranked gene within the p53 pathway group (**Fig 5C**). *Btg2* encodes B cell translocation gene 2 (BTG2), which blocks the G1-S phase transition by decreasing the levels of cyclin-dependent kinases and cyclin E and has shown to be downregulated in a variety of different cancer types (Kim *et al*, 2022). *Btg2* transcript per million (TPM) counts were shown to be only significantly downregulated in the T505A group, showing a two-fold decrease in expression compared to the MIEG3 control, wild type, R392P, and V444I (**Fig 5D**). This difference was confirmed using RT-qPCR analysis of RNA prepared from wild type and T505A ETV5-transduced cells (**Fig 5E**). Together, these findings suggest that T505A expression results in downregulation of *Btg2* gene expression, consistent with the proliferative advantage exhibited by the T505A mutation.

**Figure 5.**
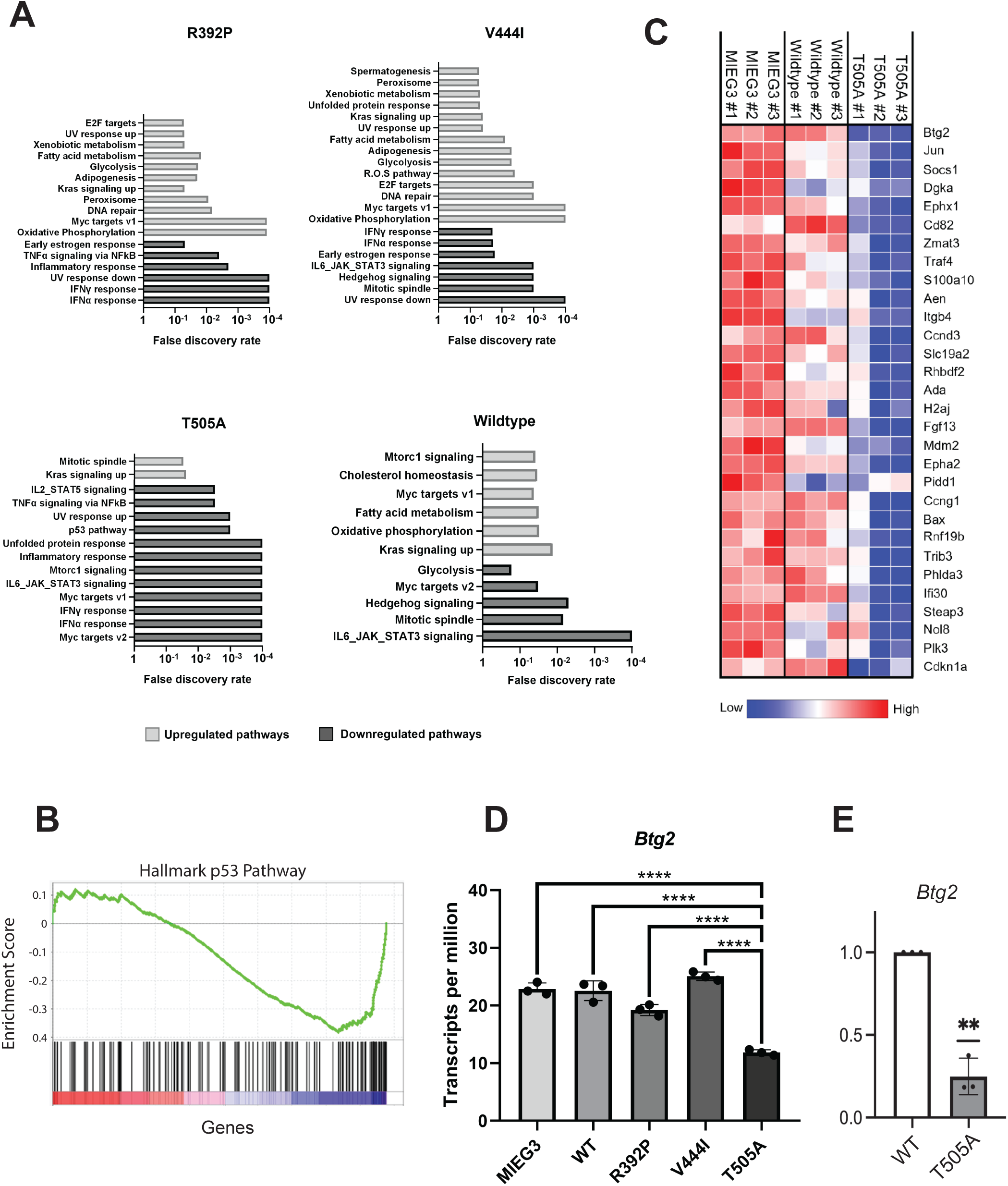
T505A ETV5 alters cell survival and proliferation pathways. **A)** Biological pathways altered in SeptMBr pre-B cells transduced with R392P (upper left), V444I (upper right), T505A (lower left), and wild type (lower right) ETV5 retroviral vectors relative to MIEG3 control-transduced cells. Upregulated pathways are indicated by light gray bars and downregulated pathways are indicated by dark gray bars. False discovery rate as determined by Gene Set Enrichment Analysis is indicated on the x-axes. **B)** Representative Gene Set Enrichment Analysis plot showing enrichment score on the y-axis and gene on the x-axis. **C)** Heat map showing relative abundances of genes in the Hallmark p53 pathway for SeptMBr cells transduced with MIEG3 control vector, wild type ETV5, or T505A ETV5. Gene names are indicated on right. **D)** Reduced number of *Btg2* mRNA transcripts per million counts (TPM) in T505A-transduced SeptMBr cells relative to cells transduced with other ETV5 variants. **E)** Reduced number of *Btg2* mRNA transcripts measured using RT-qPCR in T505A-transduced SeptMBr cells relative to cells transduced with WT ETV5. **p<0.01 by one-sample and Wilcoxon test.

## Discussion

In this study, we identified R392P, V444I, and T505A variants of ETV5 using a mouse model of pre-B-ALL. To investigate the function of these variants, we forced expression of wild type, R392P, V444I, or T505A ETV5 in an interleukin-7-dependent pre-B cell line. EMSA and luciferase assays demonstrated that the R392P and V444I variants altered DNA binding and target gene activation, while T505A and wild type ETV5 could bind DNA and activate the *Dusp6* promoter. Cell counting, proliferation, and apoptosis assays showed that R392P and V444I did not change cell count numbers, proliferation, or apoptosis. In contrast, T505A was observed to enhance cell count numbers and proliferation, as well as increase apoptosis. Using RNA-sequencing, we observed that the expression of R392P, V444I, or T505A ETV5 significantly changed gene expression, with our findings indicating that T505A downregulated the p53 pathway. Overall, these data suggest that ETV5 mutations, particularly T505A, can alter gene expression in pre-B cells to promote a leukemic phenotype.

Whole-exome sequencing revealed *Etv5* mutations were recurrent in Mb1-CreΔPB mice with 7/11 mice containing an *Etv5* mutation. Interestingly, *Etv5* mutations appeared in 6/7 ruxolitinib-treated leukemias compared to only 2/4 in control leukemias. This finding suggests that *Etv5* mutations may be positively selected by ruxolitinib; and may explain why *Etv5* mutations were not observed in our earlier studies (Lim *et al*, 2020; Batista *et al*, 2018). Ruxolitinib is a Janus Kinase inhibitor, and frequencies of *Jak1/Jak3* mutations were reduced in ruxolitinib-treated Mb1-Cre*Δ*PB mice (Mesa *et al*, 2012; Lim *et al*, 2020). *Jak1* and *Jak3* mutations are frequent in the Mb1-CreΔPB mouse model, and all leukemias in this model are interleukin-7-dependent (Batista *et al*, 2018). We speculate that suppression of interleukin-7-JAK-STAT signaling delayed leukemia progression and altered clonal evolution of leukemias by selecting for *Etv5* mutations. How ETV5 interacts with JAK-STAT signaling is unknown and will be the subject of future investigation.

Through EMSAs and luciferase assays, we found that the R392P and V444I variants, both located within the conserved ETS domain, altered DNA binding. Although no studies have specifically examined the effects of ETS domain variants on ETV5 function, our results align with findings reported for ETV1 (Cooper *et al*, 2015). In that study R391A and R394A variants in ETV1 led to a 40-fold and 1000-fold reduction in DNA-binding affinity, respectively. While R392P and V444I had weak DNA binding to a consensus oligonucleotide probe as determined by EMSA, it cannot be excluded that these variants may alter the DNA sequence specificity of ETV5. Unexpectedly, we found that forced expression of R392P and V444I ETV5 altered the expression of 578 and 515 genes, respectively, relative to wild type ETV5 (**Fig 4D, E**). The observation that these proteins alter gene expression in a manner that is distinct from one another, as well as distinct from T505A ETV5, suggests that they have altered rather than impaired DNA binding activity. ChIP-sequencing approaches will be needed to answer this question.

In contrast to R392P and V444I ETV5, EMSA and luciferase reporter experiments suggested that the T505A ETV5 variant did not affect DNA binding. This observation was expected since this amino acid is located in a C-terminal domain and not in the ETS DNA-binding domain (Laget *et al*, 1996). However, little is known about the function of this C-terminal region of ETV5. The C-terminal region of ETV5 has been shown to contain a trans-activation domain (Laget *et al*, 1996) and an autoinhibitory domain (Laget *et al*, 1996; Currie *et al*, 2017). There is no evidence for position 505 being involved in either transactivation or autoinhibition, but a role cannot be excluded. We acknowledge that M496K variants frequently coincided with T505A variants in our mouse model of leukemia (**Table 1**). We did not prioritize study of M496K since no mutations at this position have been noted in human cancers. Interestingly, most species including humans have an Alanine at position 505 of ETV5 (**Fig 1C**). Somatic A505V mutations have been noted in the COSMIC database from human intestinal and stomach cancers. We speculate that T/A505 of ETV5 plays a role in signaling by interaction with other proteins, but additional work will be needed to elucidate the function of this C-terminal region.

Forced expression of T505A ETV5 conferred a strong proliferation advantage to SeptMBr cells relative to wild type ETV5. Analysis of gene expression using RNA-seq suggested that downregulation of the p53 pathway including the gene *Btg2* was associated with proliferation of T505A ETV5-transduced cells (**Fig 5**). *Btg2* is an established tumour-suppressor gene with roles in regulating cell cycle progression and proliferation (Mao *et al*, 2015). Loss of *Btg2* has been shown to increase cell growth and proliferation in several solid tumours, including gastric, pancreatic, and lung cancers (Wei *et al*, 2012; Mao *et al*, 2015). *Btg2* has also been shown to be frequently mutated in several B cell malignancies, but its precise role in B-ALL remains unclear (Kim *et al*, 2022). Overall, we speculate that the expression of T505A in SeptMBr cells results in decreased *Btg2* expression, leading to enhanced proliferation. In future studies, the link between ETV5 and BTG2 will be investigated. Additional work will be necessary to determine what genes are direct targets of ETV5 in SeptMBr cells. ChIP-seq studies have shown that ETV5 interacts with a large number of sites in the genome relative to other transcription factors. For example, ETV5 was associated with 10,545 binding sites in type II alveolar cells (Zhang *et al*, 2017) 5,378 binding sites in Ras-transformed mammary gland epithelial cells (Arase *et al*, 2017) and 1020 binding sites in mouse embryonic stem cells (Kalkan *et al*, 2019). This large number of binding sites is associated with a significant percentage of all genes in the genome, making it challenging to use ChIP-seq data to narrow down specific target genes for this transcription factor.

A general theme that has emerged from studies on ETV5 is that it regulates and is regulated by MAP Kinase pathway of signaling in various cell types (Wei *et al*, 2023). Mitogen-activated protein kinases (MAPKs) are serine/threonine kinases that control numerous cellular processes including cell growth, proliferation, stress response, and differentiation (Suganuma & Workman, 2012). MAPK signaling has been reported to be involved in the regulation of ETV5, although the exact mechanism remains unknown (Wei *et al*, 2023). We speculate that the threonine at position 505 could be a MAPK phosphorylation site that regulates its capacity to recruit histone-modifying enzymes and remodel chromatin (Suganuma & Workman, 2012). In endothelial cells, a similar mechanism has been reported in which RAS-mediated MAPK phosphorylation of ETS proteins leads to H3 acetylation at the vascular endothelial growth factor receptor-3 (*Vegfr3*) gene (Ichise *et al*, 2012). In the other direction, ETV5 has been shown to activate the transcription of *Spry2* and *Dusp6* genes that dephosphorylate Erk-1 to downregulate MAPK signaling (Shojaee *et al*, 2015). T505A mutation may affect the ability of ETV5 to negatively regulate MAPK signaling, leading to the increased proliferation that we observed in cultured pre-B cells. More work will need to be done to elucidate the exact mechanism by which T505A ETV5 affects proliferation in pre-B-ALL.

ETV5 has been established as an oncogenic factor in a wide range of solid cancers, playing a variety of roles including cell cycling, proliferation, and drug resistance (Wei *et al*, 2023; Qi *et al*, 2020). However, despite its clear link to cancer, the role of ETV5 in leukemia is relatively unexplored. In this study, we showed that ETV5 variants can alter gene expression and provide a proliferative advantage to precursor B cells. Future studies will focus on understanding the mechanism by which ETV5 variants alter gene expression and cell behaviour.

## Materials and Methods

### Gene variant detection

Strelka (Saunders *et al*, 2012) and Varscan (Koboldt *et al*, 2013) were used to call variants from whole-exome sequencing data of 7 ruxolitinib-treated and 4 control pre-B-ALLs from Mb1-Cre*Δ*PB mice (Lim *et al*, 2020). Gene names were retrieved using BioMaRt (Durinck *et al*, 2009) and frequencies of individual variants were determined using basic functions within R Studio v4.3.2. Mutational signatures were determined using deconstructSigs (Rosenthal *et al*, 2016).

### Cloning and site directed mutagenesis

Mouse ETV5 cDNA was isolated from the MIEG3-ETV5 retroviral vector (Pham *et al*, 2014) using NotI and XhoI and ligated into the pBlueScript KS+ plasmid. Site-directed mutagenesis to generate R392P, V444I, and T505A mutants was performed on either the pBlueScript or MIEG3 plasmids using the Q5 Site-Directed Mutagenesis Kit (New England BioLabs, Boston, MA). Primers were designed using the NEBaseChanger tool (New England BioLabs). Primer sequences are listed in **Supplemental Table 1**.

### In vitro translation and electrophoretic mobility shift assay

*In vitro* translation (IVT) of WT, R392P, V444I, and T505A ETV5 proteins was performed using the TnT coupled reticulocyte lysate system (Promega, Madison, WI). For EMSA, an IRDye700-labelled consensus oligonucleotide probe was prepared and annealed (Integrated DNA Technologies, San Diego CA). Binding reactions were performed to a final volume of 20 µL containing 10 mM Tris-Cl, 1 mM EDTA, 50mM NaCl, 0.25 mM DTT, 1 µg poly dI-dC, 1 µg ETV5 probe, 0.5 µg anti-Fc or anti-ETV5 Abs, and 2 µL of IVT protein. The reactions were incubated in the dark at room temperature for 20 mins. Samples were loaded onto a 4% polyacrylamide gel at pH 7.5. Gels were pre-run at constant 30 mA for 1 hour at 4 °C, while loaded gels were run at 30 mA for 1.5 hours at 4 °C. The gel was imaged using the Odyssey CLx Imager system at 700 nm (Licor, Lincoln, NE).

### Immunoblotting

Cell lysates were prepared with 1X Laemelli buffer. Samples were denatured by boiling for 10 mins and loaded onto a 10% discontinuous polyacrylamide gel. The gel was electrophoresed at 30 mA for 1h 40 mins and proteins were transferred onto a polyvinylidene difluoride (PVDF) membrane via semi-blot transfer. The membrane was incubated overnight with blocking buffer (3:1 Intercept Blocking Buffer:PBS (Licor, Lincoln, NE) at 4 °C. Membranes incubated with 1:3000 dilution of rabbit anti-ETV5 antibodiy (ProteinTech 13011-1-AP, Rosemont IL) for 1 h, then washed 3 times with PBS/Tween. Imaging was performed using IRDye800-conjugated secondary anti-rabbit antibodies (1:20,000 dilution in PBS/Tween; Licor 926-32211). After washing 3 times with PBS/Tween, the membrane was visualized using the Odyssey CLx Imager system at 800 nm (Licor).

### Cell culture

HeLa cells and Plat-E cells (Morita *et al*, 2000) were cultured in Dulbecco’s Modified Eagle Medium (DMEM) with standard supplements (10% fetal bovine serum (Wisent, St. Bruno, Quebec), 1X penicillin/streptomycin/L-glutamine (Wisent), and 5 × 10^-5^ M β-mercaptoethanol). SeptMBr pre-B cells (Sams *et al*, 2024) were cultured at 37°C and 5% CO_2_ in Iscove’s Modified Dulbecco’s Media (IMDM) with standard supplements and 5 % IL-7 conditioned media produced by J558-IL-7 (Winkler *et al*, 1995). 38B9 cells (Alt *et al*, 1982) were cultured in RPMI-1640 (Wisent) with standard supplements.

### Retroviral transduction

Retroviral vectors were produced by transient transfection of Plat-E cells with 7.5 μg retroviral plasmid, 1 μg pCL-Eco, and 15 μL PEIpro (Polyplus, Strasbourg, France). Retroviral supernatants were used to infect SeptMBr cells by centrifugal infection at 2300 g for 2 h 30 mins at 32 °C. with 0.5 mL of virus containing 10 μg/mL polybrene and 0.5 mL 5% IL-7 IMDM.

### Flow cytometry

Retrovirally-transduced SeptMBr cells were enriched by cell sorting for green fluorescence using a BD FACSAria instrument (BD Biosciences, Franklin Lakes, NJ). Flow cytometry was performed with either a CytoFLEX S flow cytometer (Beckman Coulter, CA) or a Cytek Northern Lights cytometer (Cytek Biosciences, Brea, CA). Cell counting was performed using CountBright^TM^ beads (Thermo Fisher Scientific, Waltham, MA). Proliferation assays were performed using 24 h of culture with 0.5 μM CellTrace Violet^TM^ (Thermo Fisher Scientific, Waltham, MA). After 24 h, cells were stained with Zombie NIR viability dye (1:1000 dilution; Biolegend, San Diego, CA) and CellTrace Violet fluorescence intensity was measured using flow cytometry. Apoptosis assays were performed using the BrdU-Red TUNEL assay kit (Abcam, Cambridge, UK) with 48h of culture. Flow cytometry data was analyzed using FlowJo^TM^ v10.8.1 Software (BD Biosciences).

### Luciferase assays

Luciferase assays were performed using the Dual-Luciferase® Reporter Assay System (Promega, Madison, WI). HeLa cells were transfected with MIEG3-ETV5 vectors, DUSP6 promoter reporter, and pRL-TK control vector using PEIpro (Polyplus, Strasbourg, France). 38B9 cells were transfected using electroporation at 950 μF and 220V. Following incubation for 24 h, cells were lysed with passive lysis buffer and three rounds of freeze-thaws were performed. Luciferase activity was measured using a BioTek Cytation 5 Cell Imaging Multimode Reader (Agilent, Santa Clara, CA). Normalized luciferase values were determined by calculating the ratio of the Firefly/Renilla light units.

### Reverse transcriptase polymerase chain reaction

Total RNA was prepared from transduced SeptMBr cells using the RNeasy kit (Qiagen, Toronto, ON) and cDNA was synthesized using the iScript cDNA Synthesis kit (Bio-Rad, Missisauga, ON). Forward and reverse primers were designed using the PrimerQuest Tool (Integrated DNA Technologies, San Diego CA, United States). RT-qPCR was performed using the QuantStudio 5 PCR System (Thermo Fisher Scientific, Mississauga, ON). Gene expression was measured in triplicate using the ΔΔCT method.

### RNA-sequencing

RNA sequencing was performed on total RNA prepared from transduced SeptMBr cells using a polyA-enriched RNA library prep Kit and Illumina NovaSeq-6000 PE 100-25M reads (Illumina, San Diego, CA) FASTQ files were processed using the following pipeline with Galaxy (https://usegalaxy.org): (1) Trimmomatic (Bolger *et al*, 2014; Patro *et al*, 2017) to trim sequences, (2) generation of GRCm38-aligned BAM files using HiSat2 (Kim *et al*, 2019), (3) Salmon to calculate transcript counts (Patro *et al*, 2017), and (4) DEseq2 (Love *et al*, 2014) to determine differentially expressed genes. Differentially expressed genes were identified as genes with a padj < 0.05 and log2FoldChange > 0.5. Gene set enrichment analysis (Subramanian *et al*, 2005) (GSEA) was performed on all hallmark pathways identified in the GSEA pathway library (mh.all.v2023.2). Significantly enriched pathways were identified using a false discovery rate q-value < 0.05. Heat maps and volcano plots were generated using the heatmap function in RStudio v4.3.2.

### Statistics

Statistical analysis was performed using Prism 10 (Graphpad, San Diego, CA) based on independent biological replicate experiments. Individual statistical tests are named in the figure legends.

## Data Availability

RNA sequencing data is available as BAM files (GRCm38) from the Sequence Read Archive, BioProject Accession PRJNA1200055.

## Supporting information

Supplemental Gene List

Supplemental Gene List

Supplemental Gene List

Supplemental Gene List

Supplemental Variant List

Supplemental Variant List

## Acknowledgements

We thank Dr. Mark Kaplan (Indiana University) for providing us with the MIEG3 and MIEG3-ETV5 retroviral vectors. We thank Genome Quebec (Montreal, Quebec) for assistance with RNA-sequencing. We thank Kristin Chadwick and the London Regional Flow Cytometry Core Facility for assistance with cell sorting and analysis. This work was supported by Canadian Institutes of Health Research grant 168995.

## Author contributions

**Rodney P. DeKoter:** Conceptulization; funding acquisition; methodology; writing – review and editing; project administration.

**Joshua Yi:** Conceptualization; methodology; writing – original draft.

**Michael Yu:** Methodology; writing – original draft.

**Michaela L. Dowling:** Methodology; writing – original draft.

**Allanna C. E. Mackenzie:** Methodology.

**James Iansavitchous:** Methodology.

