## Supplemental Data for "T505A variant of ETV5 promotes proliferation of precursor B cells in a mouse model of acute lymphoblastic leukemia"

**Appendix – Yi et al.**

**Supplemental Data**

Two Excel spreadsheets are provided containing lists of variants called by Strelka and Varscan:

Strelka Filtered Genes (0.10, MSSS).xlsx
Varscan Filtered Genes (0.10, MSSS).xlsx

Four tab-separated files are provided containing lists of differentially expressed genes identified using DESEQ2:

Genelist_MIEG3_VS_WT.tsv
Genelist_MIEG3_VS_V444I.tsv
Genelist_MIEG3_VS_R392P.tsv
Genelist_MIEG3_VS_T505A.tsv

**Table S1. Primer Sequences**

| Purpose | Primer | Sequence (5' to 3')[a] |
|---|---|---|
| ETV5 site directed mutagenesis Arg392Pro | Forward | TGGACTGGCC**C**AGGCATGGAA |
| | Reverse | GGCGATGAAGTGGGCATTG |
| ETV5 site directed mutagenesis Val444Ile | Forward | GGAACGCTAC**A**TCTACAAATTTGTG |
| | Reverse | CCAGCCACCTTCTGCATG |
| ETV5 site directed mutagenesis Thr505Ala | Forward | CCTCCCCTAC**G**CCGAAGGCTT |
| | Reverse | CTGCTGCAGCGATCCATG |
| Etv5 Sequencing 906-924 | Forward | CTCAGAAGTGCCTAACTGC |
| Etv5 Sequencing 1395-1413 | Forward | GTTCCTGAAGGCAGAATCC |
| Etv5 Sequencing 133-115 | Reverse | CTTCAGAATCGTGAGCCAG |
| Etv5 Sequencing Reverse 2 623-604 | Reverse | GGCATCATCTTTGGCATTTG |
| ECM Probe for EMSA | Forward | 5IRD700/AAA**CCGGAA**GTCTG |
| | Reverse | 5IRD700/CAGACTTCCGGTTT |
| Mutated ECM Probe for EMSA | Forward | 5IRD700/AAA**CCGCAA**GTCTG |
| | Reverse | 5IRD700/CAGACTTGCGGTTT |
| DUSP6 promoter cloning | Forward | GCTTGGATCTAGCGTTAAAGAGAATTA |
| | Reverse | CAATGAATCTCTCTCTCTGAATGAAGG |

| | | |
|---|---|---|
| Btg2 RT-qPCR | Forward | CGCACTGACCGATCATTACAA |
| | Reverse | GGGTCCATCTTGTGGTTGATAC |

[a] Underlined and bolded nucleotides indicate incorporated single-nucleotide polymorphism.

**Table S2. Identities of Mice Included in Ruxolitinib Chow Experiment**

| Mouse ID | Ruxolitinib Chow | Sex | Age at euthanasia (weeks) | Strelka Variants (>0.1 MSSS) | Varscan Variants (>0.1 MSSS) |
|---|---|---|---|---|---|
| 229 | None | F | 14.3 | 73 | 336 |
| 258 | None | M | 17.4 | 79 | 953 |
| 262 | None | M | 17.4 | 356 | 1485 |
| 484 | None | F | 15.3 | 105 | 1443 |
| 224 | 4-8 wk | F | 58.1 | 87 | 719 |
| 226 | 4-8 wk | F | 27.3 | 84 | 245 |
| 261 | 4-8 wk | F | 22.7 | 155 | 1637 |
| 307 | 4-8 wk | F | 27.7 | 65 | 724 |
| 310 | 4-8 wk | F | 27.7 | 31 | 428 |
| 339 | 4-8 wk | F | 51.9 | 165 | 1333 |
| 486 | 4-8 wk | M | 40.0 | 204 | 1863 |

**Table S3. Identification of Gene Variants.**

| Gene | Frequency (per 11 mice) | Variant Caller | HGNC symbol | Cancer Gene Census |
|---|---|---|---|---|
| Gm9573 | 11 | Varscan | MUC21 | No |
| Zfp473 | 11 | Varscan | ZNF473 | No |
| Fan1 | 10 | Varscan | FAN1 | No |
| Stat2 | 10 | Varscan | STAT2 | No |
| Tchh | 10 | Varscan | TCHH | No |
| Ttn | 10 | Varscan | TTN | No |
| Herc2 | 9 | Varscan | HERC2 | No |
| 4932438A | 8 | Varscan | KIAA1109 | No |
| Dst | 8 | Varscan | DST | No |
| Etv5 | 8 | Varscan | ETV5 | Tier 1 |
| Macf1 | 8 | Varscan | MACF1 | No |
| Mrgprx1 | 8 | Varscan | MRGPRX1 | No |
| Ppp1r15a | 8 | Varscan | PPP1R15A | No |
| Vps13a | 8 | Varscan | VPS13A | No |
| Ascc3 | 7 | Varscan | ASCC3 | No |
| Atm | 7 | Varscan | ATM | Tier 1 |
| Ccdc114 | 7 | Varscan | CCDC114 | No |
| Csmd1 | 7 | Varscan | CSMD1 | No |
| Dux | 7 | Varscan | DUX1 | No |

**Table S4**. **Near-Cognate Translation Initiation Site Prediction Analysis on Wildtype ETV5 cDNA**

| Nucleotide Position | Predicted MW[a] | Kozak Similarity Score[b] |
|---|---|---|
| 634 | 35 kDa | $\geq 0.6 - \leq 0.7$ |
| 754 | 30 kDa | $\geq 0.7 - \leq 0.8$ |
| 847 | 27 kDa | $\geq 0.6 - \leq 0.7$ |
| 897 | 25 kDa | $\geq 0.6 - \leq 0.7$ |

[a] Molecular weights were predicted using the ExPASy software.

[b] Kozak similarity scores were predicted using TIS predictor software; higher scores indicate higher similarities to a Kozak sequence.

**Figure S1. Variant calling and mutational signature analysis of whole-exome sequencing of 11 pre-B-ALLs from Mb1-CreDPB mice. A)** Schematic of workflow for variant calling. **B)** Number of variants called using Strelka or Varscan algorithms. Number of variants called using Strelka (filled circles) are connected to number of variants called from same leukemia using Varscan (open circles. **C)** Representative example of mutational signature analysis using deConstructSigs. Two representative leukemias are shown: 229 (control, top) and 224 (ruxolitinib-treated, bottom). Mutation types are indicated in colour using legend on right. Dinucleotide context of mutations is shown in x-axis. **D)** The ratio of C>A to C>T variants (y-axis) is higher for low variant allele frequencies compared to high variant allele frequencies (x-axis). ***$p<0.001$ using t-test.

**Figure S2. ETV5 protein interaction with consensus DNA oligonucleotide probe is disrupted by single base change. A)** DNA sequence of consensus DNA oligonucleotide probe for ETV5 (top) and DNA sequence of consensus DNA oligonucleotide probe with one base change (C -> C). **B)** Electrophoretic mobility shift assay for ETV5 variants using unmutated and mutated probes. In vitro translated proteins (top row) were incubated with the indicated probes (second from top row). Location of shifted and free probe is indicated on right.

**Figure S3. Activation of a DUSP6 promoter luciferase reporter by the ETV5 C-terminal transactivation domain. A)** The DUSP6 promoter is active in 38B9 pre-B cells. The DUSP6 promoter luciferase reporter was transiently transfected into 38B9 mouse pre-B cells by electroporation. Relative light units (y-axis) were determined as ratio of light units from luciferase reporter vector to pRL-TK Renilla luciferase control reporter. X-axis indicates positive control (pGL3-control), negative control (pGL3-basic), and test (pGL3-DUSP6). **B)** The DUSP6 promoter is active in HeLa cells. The DUSP6 promoter luciferase reporter was transiently transfected into HeLa cells using PEIpro reagent. Relative light units (y-axis) were determined as ratio of light units from luciferase reporter vector to pRL-TK Renilla luciferase control reporter. X-axis indicates positive control (pGL3-control), negative control (pGL3-basic), and experiment (pGL3-DUSP6). **C)** Schematic of the DUSP6 promoter luciferase reporter vector (top) and the MIEG3-ETV5 C-terminal expression vector (bottom). **D)** Activation of the DUSP6 promoter by the wild-type (wildtype) or T505A mutated C-terminal region of ETV5. Relative light units (y-axis) were determined as ratio of light units from luciferase reporter vector to pRL-TK Renilla luciferase control reporter. X-axis indicates plasmid co-transfected with DUSP6 promoter reporter vector: MIEG3 (empty vector) MIEG2-WT ETV5 C-term (wildtype) or MIEG-T505A C-term (T505A).

**Figure S4. Retroviral transduction of SeptMBr Pre-B cells with ETV5 variants. A)** Flow cytometric analysis of SeptMBr pre-B cells following enrichment by cell sorting based on green fluorescence. Legend indicates non-transduced (red) or transduced (blue). Frequencies of cells in green fluorescence gate is indicated on histogram. **B)** Relative expression of ETV5 mRNA transcripts using RT-qPCR analysis. Y-axis indicated relative gene expression. X-axis indicates retroviral transduction condition. **$p<0.01$ by one-way ANOVA with Tukey's test. **C)** Anti-ETV5 immunoblot. Top panel shows anti-ETV5 immunoblot, lower panel shows anti-beta-actin immunoblot.

**Figure S5. Unique and shared differentially expressed genes comparing all ETV5 variant groups.** Shown is Venn diagram with numbers indicating numbers of up- or down-regulated genes unique to each ETV5 variant group, or in common between all four groups.

### Figure S1

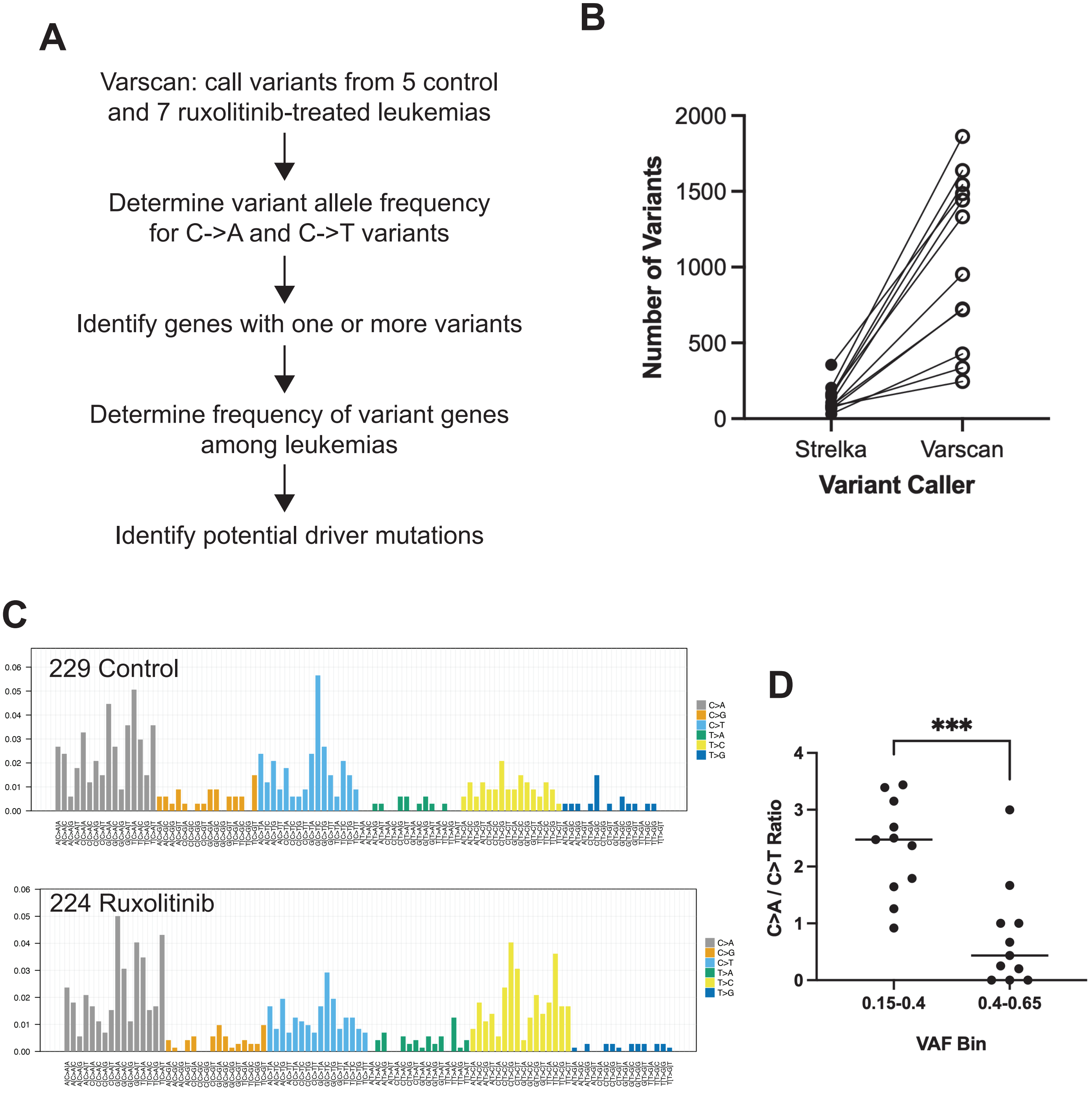
A
Varscan: call variants from 5 control and 7 ruxolitinib-treated leukemias
Determine variant allele frequency for C->A and C->T variants
Identify genes with one or more variants
Determine frequency of variant genes among leukemias
Identify potential driver mutations
B
Number of Variants
0
500
1000
1500
2000
Strelka
Varscan
Variant Caller
C
229 Control
224 Ruxolitinib
0.00
0.01
0.02
0.03
0.04
0.05
0.06
C>A
C>G
C>T
T>A
T>C
T>G
D
***
C>A / C>T Ratio
0
1
2
3
4
0.15-0.4
0.4-0.65
VAF Bin

#### Figure S2

**A**

Probe: 5’/IRD700/AAACCGGAAGTCTG/3’

Mutated Probe: 5’/IRD700/AAACCG<u>C</u>AAGTCTG/3’

**B**

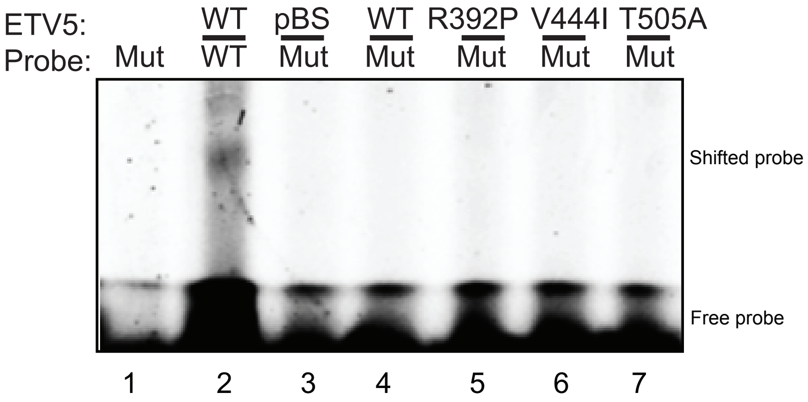

### Figure S3

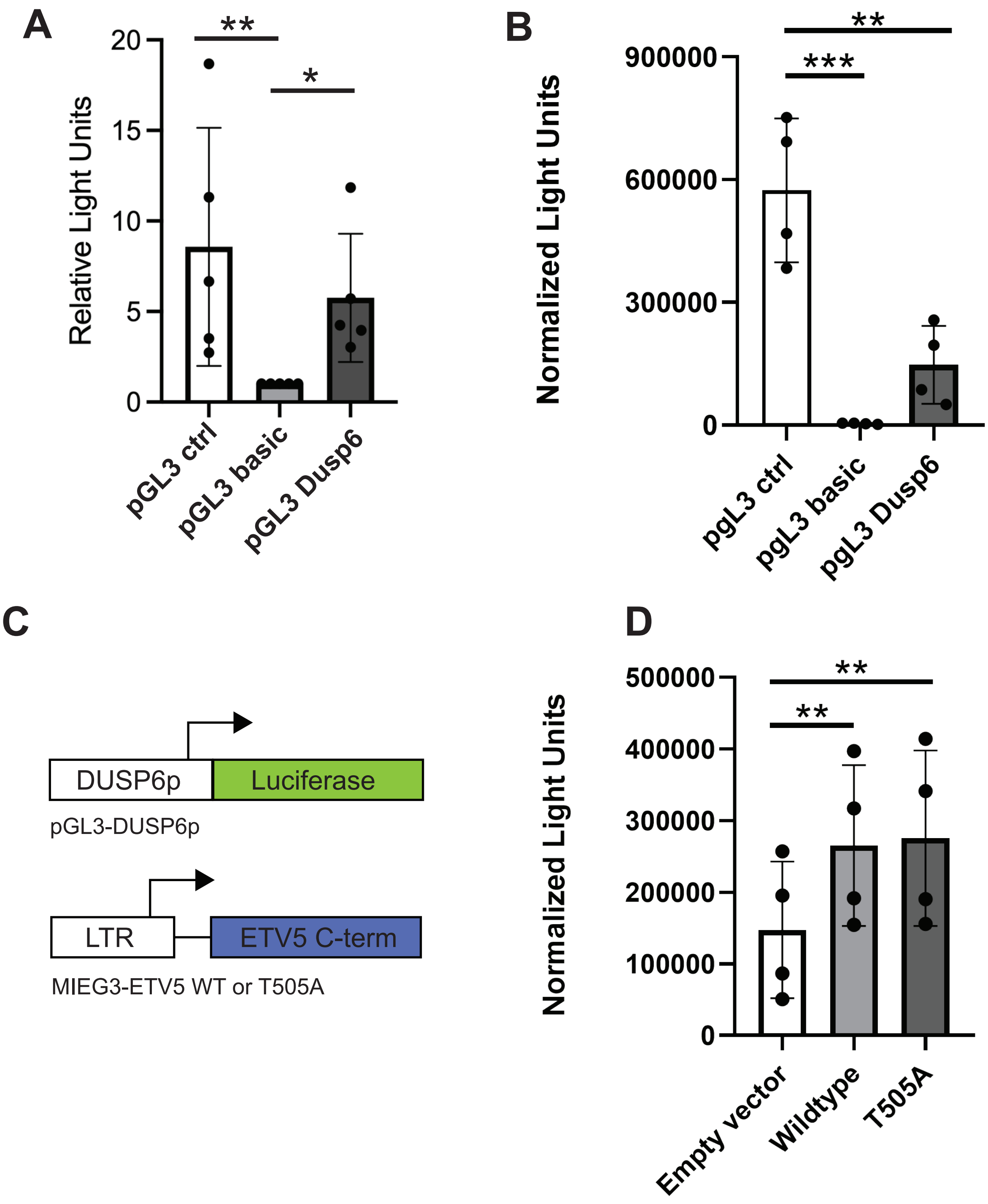

A
Relative Light Units
20
15
10
5
0
**
*
pGL3 ctrl
pGL3 basic
pGL3 Dusp6
B
Normalized Light Units
900000
600000
300000
0
**
***
pgL3 ctrl
pgL3 basic
pgL3 Dusp6
C
DUSP6p
Luciferase
pGL3-DUSP6p
LTR
ETV5 C-term
MIEG3-ETV5 WT or T505A
D
Normalized Light Units
500000
400000
300000
200000
100000
0
**
**
Empty vector
Wildtype
T505A

### Figure S4

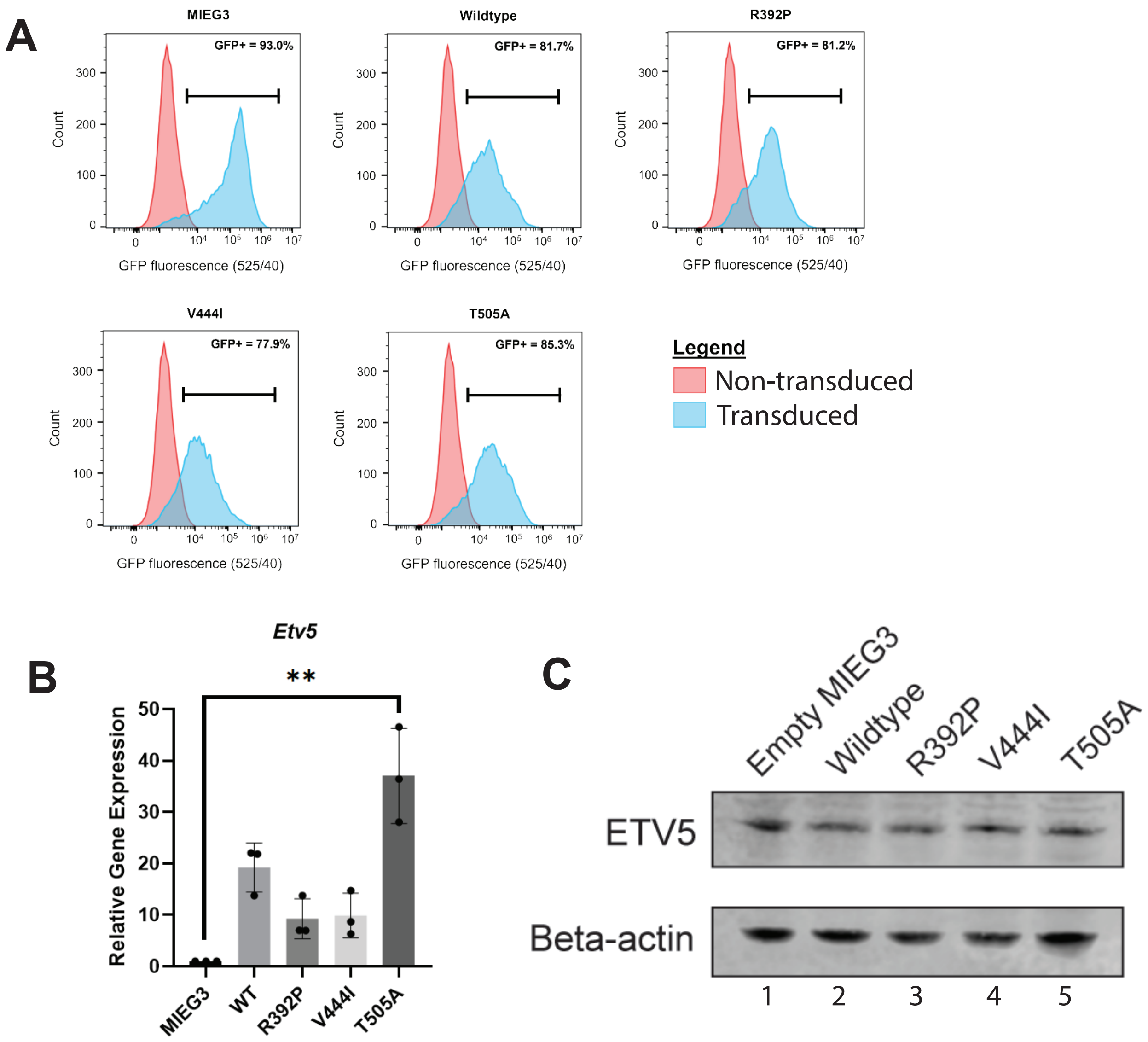
A
MIEG3
GFP+ = 93.0%
Wildtype
GFP+ = 81.7%
R392P
GFP+ = 81.2%
V444I
GFP+ = 77.9%
T505A
GFP+ = 85.3%
Count
GFP fluorescence (525/40)
Legend
Non-transduced
Transduced
B
Etv5
**
Relative Gene Expression
MIEG3
WT
R392P
V444I
T505A
C
Empty MIEG3
Wildtype
R392P
V444I
T505A
ETV5
Beta-actin
1
2
3
4
5

**Figure S5**

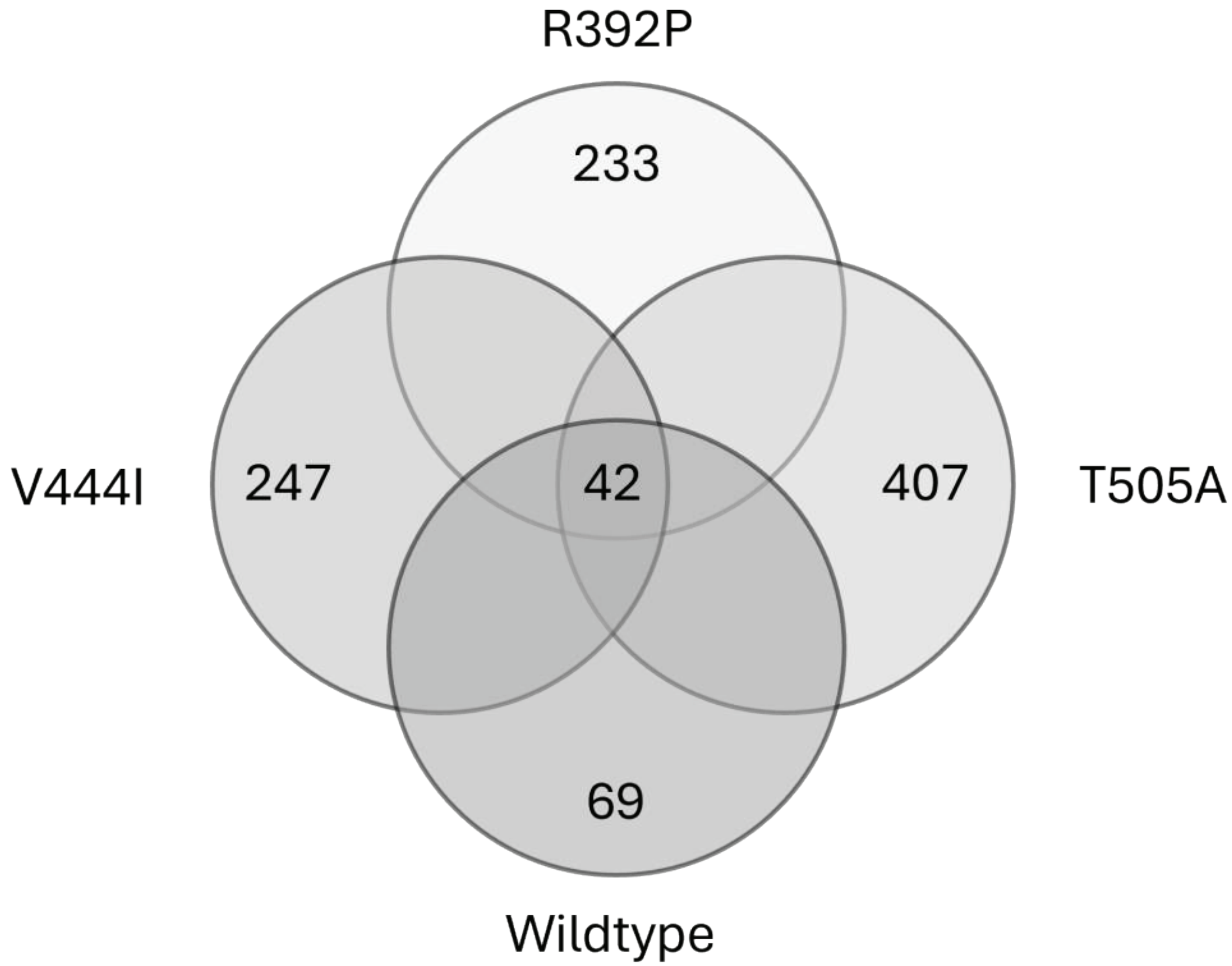
